# Neurodevelopmental shifts in learned value transfer on cognitive control during adolescence

**DOI:** 10.1101/801126

**Authors:** Catherine Insel, Mia Charifson, Leah H. Somerville

**Affiliations:** Department of Psychology and Center for Brain Science, Harvard University, Cambridge, MA, USA

**Keywords:** adolescence, cognitive control, learning, value, striatum, prefrontal cortex

## Abstract

Value-associated cues in the environment often enhance subsequent goal-directed behaviors in adults, a phenomenon supported by integration of motivational and cognitive neural systems. Given the interactions among these systems change throughout adolescence, we tested when beneficial effects of value associations on subsequent cognitive control performance emerge during adolescence. Participants (N=81) aged 13-20 completed a reinforcement learning task with four cue-incentive pairings that could yield high gain, low gain, high loss, or low loss outcomes. Next, participants completed a Go/NoGo task during fMRI where the NoGo targets comprised the previously learned cues, which tested how prior value associations influence cognitive control performance. Improved accuracy for previously learned high gain relative to low gain cues emerged with age. Older adolescents exhibited enhanced recruitment of the dorsal striatum and ventrolateral prefrontal cortex during cognitive control execution to previously learned high gain relative to low gain cues. Older adolescents also expressed increased coupling between the dorsal striatum and dorsolateral prefrontal cortex for high gain cues, whereas younger adolescents expressed increased coupling between the striatum and ventromedial prefrontal cortex. These findings reveal that learned high value cue-incentive associations enhance cognitive control in late adolescence in parallel with value-selective recruitment of corticostriatal systems.

## Introduction

Adolescence is a time of life when learning is paramount. As adolescents navigate their worlds, they encounter many new situations, both good and bad, and they must use the outcomes of these situations to guide their future decisions and actions. For example, imagine an adolescent enters the cafeteria, looking for a place to eat lunch. They must first learn from positive and negative feedback to determine where it is appropriate to sit and who is willing to sit with them. Following these encounters, the learned associations from these initial interactions, both good and bad, may generalize across contexts. Therefore, when encountering these cues in new environments, their learning history will orient their attention towards high value cues. For instance, upon seeing peers from prior interactions in the hallway or classroom, they will focus attention towards a friendly classmate. As such, learning from feedback can create longstanding consequences for goal-directed behavior. However, it remains unclear when and how adolescents transfer these learned associations to new contexts when immediate feedback is no longer provided. Given that the brain systems that integrate information about incentive value and action selection continue to mature throughout adolescence (Insel et al., 2017), here we tested how the ability to transfer value associations across contexts to guide goal-directed behavior emerges with age during adolescence.

Adolescence is a period of the lifespan when brain systems that support cognition and motivation continue to mature to guide the emergence of successful goal-directed behavior (Davidow et al., 2018a; Somerville & Casey, 2010), and adolescents continue to refine their cognitive abilities with age (Casey, 2015; Paus, 2005). The ability to implement cognitive control, a mental process that allows individuals to select contextually appropriate behavior to pursue superordinate goals (Miller & Cohen, 2001), continues to improve through late adolescence and into early adulthood (Crone & Steinbeis, 2017; Luna et al., 2010). This protracted development of cognitive control is related to ongoing structural and functional maturation of the prefrontal cortex (Baum et al., 2017; Satterthwaite et al., 2013; Tang et al., 2017). Moreover, strategic improvements in cognitive control emerge with age alongside enhanced functional connectivity between the striatum and prefrontal cortex (Vink et al., 2014).

While the majority of prior developmental research has tested cognitive control to neutral cues, a growing body of work investigates how cues denoting motivational value interact with the execution of cognitive control differentially with age (Cohen et al., 2016; Davidow et al., 2018b; Hare et al., 2008; Somerville et al., 2011). Findings from some of these studies suggest that adolescents exhibit behavioral disruptions when exerting control to actively appetitive cues relative to earlier and later ages, and this adolescent-unique behavioral expression is linked to age-related differences in recruitment of neural systems implicated in reward processing and cognitive control (Casey, 2015). These prior findings in adolescents may reflect Pavlovian-instrumental transfer, whereby the biases to approach reward related cues increase instrumental responding (Cartoni et al., 2016; Talmi et al., 2008). Therefore, if a learned reward association is especially strong, it may be even more difficult to successfully engage response inhibition when the cue is presented, such as in the Go/NoGo tasks described above, which require individuals to stop prepotent responses to reward-related cues. Therefore, reward associations may hinder subsequent control performance when these associations are yoked to motor actions.

However, in prior work on adults, there are situations when prior reward history can yield beneficial effects and enhance subsequent cognitive control performance. In adults, cues that are otherwise neutral but hold value based on learning history can modulate attention and facilitate behavior, which can in turn bolster goal-relevant action selection as well as enhance the salience and detection of motivated cues in the environment (Corbetta & Shulman, 2002). Prior work in adults has examined how previous value associations influence cognitive control by first training associations between visual cues and incentive outcomes, and then testing how individuals perform in a subsequent cognitive task when a previously-valued cue is present (Krebs & Woldorff, 2017). Typically, the presence of a previously-valued cue yields a performance benefit, even when the subsequent task is no longer incentivized. For example, prior work using a visual search task found that cues with prior reward associations enhanced attention when incentives were no longer at stake (Anderson et al., 2011a, 2011b). Further, prior research has found that when low and high value associations are first trained, and then these previously rewarded cues are inserted in a subsequent working memory task, working memory improves for cues that once predicted high value outcomes (Gong & Li, 2014). Together, this prior work in adults suggests that the presence of previously valued cues can orient attention and facilitate goal directed behavior.

While previously-incentivized cues can benefit adult cognitive performance, it remains unclear how adolescent cognitive control is impacted by prior reinforcement history. A growing body of work has tested adolescent cognitive performance when actions are actively incentivized, and converging evidence suggests that adolescents are less likely than adults to enhance performance according to the value of their goals (Castel et al., 2011; Stormer et al., 2014), with the ability to do so emerging in late adolescence (Insel et al., 2017). This late developing ability to integrate value goals with control performance may be related to ongoing development of corticostriatal circuitry (Davidow et al., 2018a). For example, when sustained attention is rewarded, recruitment of the dorsal striatum and prefrontal cortex increases with age from childhood to adulthood during rewarded compared to neutral cognitive performance (Smith et al., 2011). Further, when cognitive control is incentivized, older adolescents and young adults selectively increase the recruitment of the striatum and prefrontal cortex during the execution of cognitive control when high stakes outcomes are at play, but younger adolescents do not (Insel et al., 2017).

However, it remains unclear whether this age-related pattern in value-selective behavior and corticostriatal recruitment persists when previously-valued cues are presented during cognitive control execution, but performance is no longer incentivized. Specifically, we aimed to test whether value associations acquired through learning persist beyond the learning environment to modulate subsequent performance. If value history confers a persistent beneficial effect on performance, this would reveal that learned value associations can transfer across cognitive domains and modulate attentional control. Moreover, by testing the transfer of behavior when performance is no long incentivized, we can isolate how learned cue-incentive associations can confer lasting benefits to behavior.

Specifically, this study tested whether the valence (gain vs. loss) and magnitude (high vs. low) of learned incentives differentially influence subsequent control performance to resolve two theoretical accounts of value-related control performance. On the one hand, if gain learning enhances approach behavior and loss learning enhances avoidance behavior, previously learned gain cues should dampen response inhibition whereas previously learned loss cues should enhance response inhibition, a profile which has been observed in prior studies testing adults (Guitart-Masip et al., 2014). On the other hand, if gain learning sharpens visual attention and discrimination more than loss learning, we would expect to see improved cognitive control to previous gain cues, in line with prior work (Krebs & Woldorff, 2017). Finally, the magnitude manipulation allows us to test not just how valence-associations influence control performance, but whether these helping or hindering effects scale with the magnitude of learned incentive value.

To do so, we first trained low and high gain and loss value associations with specific cues during reinforcement learning, and then tested the ability to withhold responses to these previously learned cues during a Go/NoGo task when performance is no longer incentivized. Behavioral analyses identified age-related patterns in how previously learned cue-incentive associations interact with subsequent cognitive control performance. We specifically queried whether prior low and high stakes value differentially bias subsequent performance for cues previously associated with gains versus losses. Neuroimaging analyses examined age-related differences in functional recruitment during cognitive control that, with age, varied as a function of prior stakes value.

## Methods

### Participants

84 male and female participants age 13 to 20 participated in this study (mean age = 16.97, standard deviation = 2.47). Participants were recruited from the greater Boston area and screened for current psychiatric or neurological illness, history of learning disabilities, proficiency of the English language, and had no lifetime use of psychotropic medication. Before study participation, participants and their legal guardians (for minors) provided written assent and consent under the protocol approved by the Committee for Use of Human Subjects at Harvard University. The total sample included 81 participants for the behavioral analysis and 74 participants for the fMRI analysis.

Two participants were excluded from behavioral analysis due to technical issues with data collection, and one participant was excluded for responding to fewer than 50% of trials during the experimental task. Two participants did not complete the fMRI scan and performed the task behaviorally outside of the scanner. Five participants were excluded from fMRI analysis due to excessive motion (see the section *FMRI Data Processing and Quality Assessment* below for details).

### Reinforcement learning task

Participants completed a reinforcement learning task (Figure 1A) in which they repeatedly viewed pairs of fractals, one of which was the optimal choice for accruing financial earnings. Participants selected among the pairs and received trial-by-trial probabilistic feedback indicating monetary payouts, and participants could use this feedback to learn to orient their subsequent choices toward the optimal fractal.

**Figure 1.**
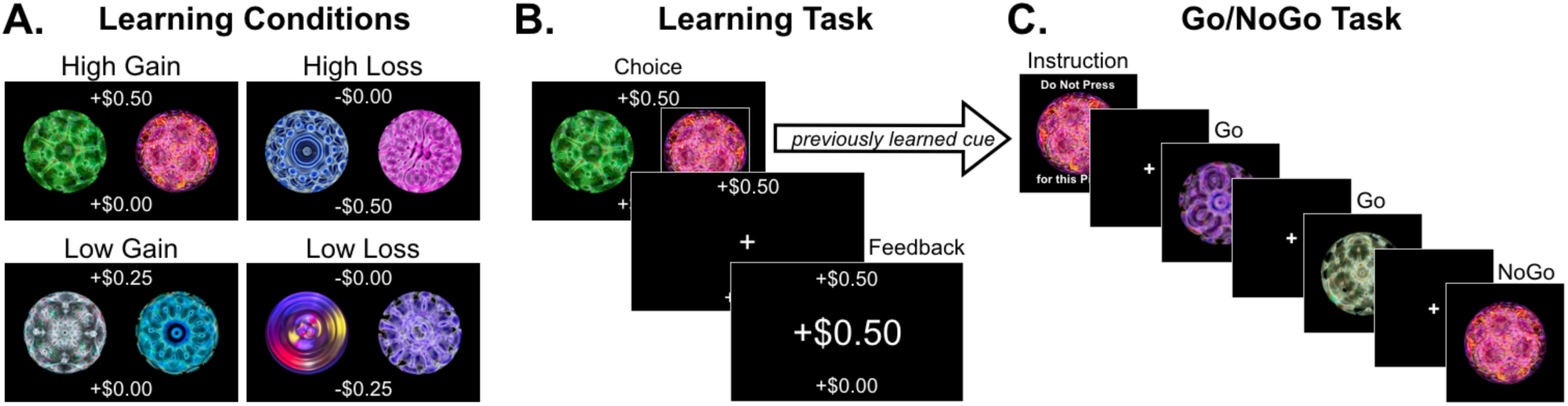
Experimental Tasks. **A) Learning Conditions**. Learning task conditions consisted of high gain, low gain, high loss, and low loss. The monetary outcomes associated with the optimal and suboptimal choices were displayed at the time of choice on the top and bottom of the screen. **B) Example learning trial.** Participants chose one fractal and then received probabilistic monetary feedback. Following an optimal choice, participants earned favorable feedback on 75% of trials. **C) Post-Learning Go/NoGo Task.** Following the learning task, participants completed a Go/NoGo Task in which the NoGo cues comprised the previously learned cues associated that were most often associated with winning or losing money. There were no incentives for performance during the Go/NoGo Task.

**Figure 2.**
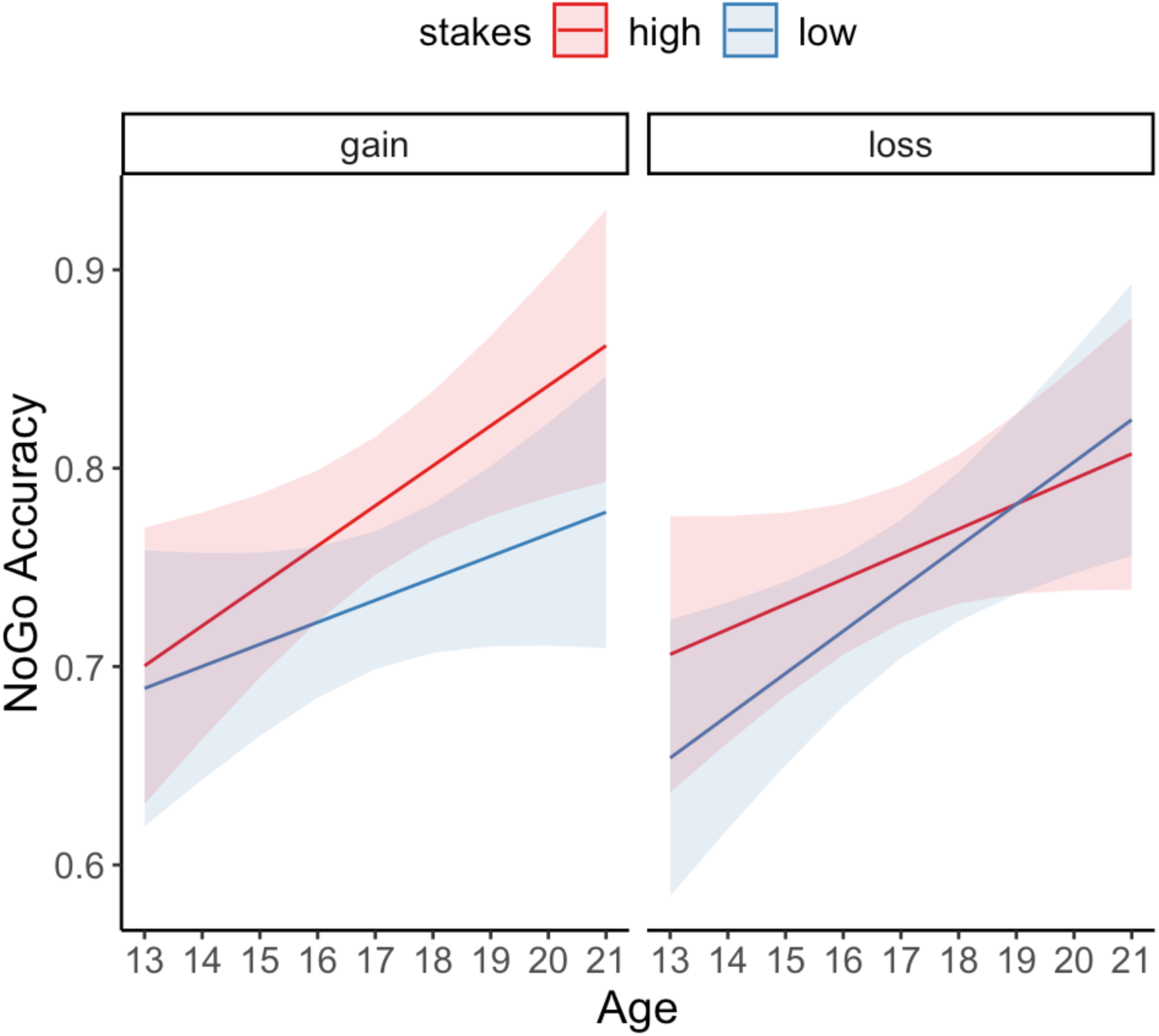
Significant interaction between age, stakes, and valence on post-learning NoGo accuracy. NoGo accuracy is plotted for cues previously associated with high stakes (+50¢ or - 50¢, plotted in red) and low stakes (+25¢ or -25¢, plotted in blue) incentives within gain (left panel) and loss (left panel) conditions across age. Lines represent linear age fits and shaded bands represent between-subjects 95% confidence intervals.

The task included four conditions consisting of distinct fractal pairings, where each pair of fractals was associated with different monetary stakes and valence outcomes: high gain (+$0.50/+$0.00¢), low gain (+$0.25/+$0.00), high loss (-$0.50/-$0.00), and low loss (-$0.25/-$0.00). The potential outcomes at stake were displayed at the time of choice at the top and bottom of the screen to minimize working memory demands. Participants were instructed that their earnings would be paid out at the end of the study. In the event that a participant ended up with a net loss, no money was taken away.

For each trial, two fractal images appeared on the screen, and participants chose one fractal by pressing the index finger of the right hand to choose the fractal on the left side of the screen or the middle finger to choose the fractal on the right side. Once a choice was made, a box appeared on the screen surrounding the chosen fractal. Monetary feedback was then displayed to indicate the outcome of the choice. For gain trials, choosing the optimal fractal resulted in a monetary gain, whereas for loss trials choosing the optimal fractal resulted in no losses. Feedback was probabilistic such that the optimal fractal yielded the favorable outcome for 75% of trials, whereas suboptimal fractal yielded unfavorable outcomes on 75% of trials. If a participant failed to choose a fractal in the time window (missed trial), they received the unfavorable outcome.

Participants completed 240 learning trials, with 60 trials per incentive condition. Trials were split evenly across four blocks, with 60 trials per block and 15 trials of each condition intermixed throughout the block. Condition order was pseudo-randomized within and across runs. Fractal pairs were displayed for 3 seconds; when the participant made a choice, a selection box appeared to highlight the chosen fractal. Next a jittered fixation was displayed (1.5-3 seconds). Finally, monetary feedback was displayed for 1 second. Fractal display was counterbalanced across trials (left versus right side presentation). Fractal condition assignment was counterbalanced across participants.

### Post-learning Go/NoGo task

Following the reinforcement learning task, participants completed a Go/NoGo task during functional neuroimaging. In this task, participants first viewed an instruction screen, which displayed the NoGo fractal image for the next block, and participants were instructed to not press a button when this image appeared during the forthcoming block of trials. Next, a series of 18 fractal images were displayed, consisting of Go targets and NoGo targets. Participants were instructed to press the index finger of their right hand as quickly as possible for Go targets but to withhold responding for NoGo targets. Go targets consisted of novel fractal images that the participants had not encountered in the prior task. NoGo stimuli consisted of the fractals displayed during the learning task that were previously associated with the greatest chance of winning or losing money (e.g. low gain optimal choice image, high gain optimal choice image, low loss suboptimal choice image, high loss suboptimal choice image). Importantly, participants were not aware during learning that these images would be carried forward to a subsequent task, and Go/NoGo performance was not incentivized.

Participants completed 12 blocks of 18 trials during the Go/NoGo task. For each block of trials, participants first viewed an instruction screen for 2 seconds that displayed the NoGo image for the upcoming block of trials. Next, a series of 18 target trials were presented for 500 milliseconds each followed by a jittered fixation cross (1.5-3 seconds). At the end of the block, a fixation cross was displayed for 3 seconds. Target trials consisted of 66% Go trials and 33% NoGo trials. The prepotency of Go trials preceding a NoGo ranged from 0 to 4 Go trials, and prepotency order was pseudorandomized across blocks within each condition and balanced across conditions.

### Practice Go/NoGo task

Before the fMRI session, participants completed a practice Go/NoGo task with a different set of fractal images that had no prior learning history and were not included in the post-learning Go/NoGo stimuli set. The practice task consisted of 24 NoGo and 48 Go trials, which were presented across 4 blocks of 18 trials each. Each practice block contained a unique NoGo image. The task presentation parameters, such as NoGo frequency and Go prepotency, were identical to the Post-Learning Go/NoGo.

### Task ratings to assess subjective value of monetary outcomes

To measure individual differences in subjective value of the monetary outcomes in the learning task, participants made subjective pleasantness and arousal ratings for how they felt when earning each possible monetary outcome: +$0.50 and +$0.00 in the high gain condition, +$0.25 and +$0.00 in the low gain condition, -$0.50 and -$0.00 in the high loss condition, and - $0.25 and -$0.00 in the low loss condition. These ratings served as a manipulation check to verify that participants differentially valued the low and high gain and loss outcomes. Ratings were collected using the Self-Assessment Manikin scales for pleasantness (how good or bad did that make you feel) and arousal (how intense was that feeling) (Lang, 1980) with numerical anchors. Pleasantness ratings were made on a scale from 1 (unpleasant) to 9 (pleasant). Arousal ratings were made on a scale from 1 (low arousal) to 9 (high arousal). These ratings were not collected from four participants due to technical issues with data collection.

### Behavioral analysis

Learning Performance: Learning performance was calculated as the proportion of trials on which the participant chose the optimal fractal in the late phase of the task (trials 180-240). The optimal fractal was operationalized for each pair of trials as the one that delivered the most favorable outcome the majority of the time. In the gain conditions, the most favorable outcomes resulted in monetary reward and in the loss conditions the most favorable outcomes resulted in no loss (lose 0¢). Optimal choice performance was analyzed with a mixed-effects model to test for fixed effects of stakes (low/high), valence (gain/loss), continuous linear age, and all interaction terms. The model included a random effect for subject to account for repeated measures.

Go/NoGo Performance: Behavioral analyses assessed Go/NoGo performance to determine whether prior value associations transferred across tasks and influenced cognitive control to previously learned cues. The primary dependent variable was NoGo accuracy, which was calculated as the proportion of trials on which participants successfully withheld a button response to NoGo cues. NoGo trials comprised cues that had been previously associated with low stakes and high stakes gain and loss outcomes in a prior learning task. We conducted a mixed-effects model to test for main effects of stakes (low/high) and valence (gain/loss) of the previously learned cues and their interaction on NoGo accuracy. This model included fixed effects for stakes, valence, and a stakes by valence interaction and included a random effect for subject to account for repeated measures. The goal of this analysis was to establish whether previously learned cue-incentive associations transferred across tasks to influence subsequent NoGo performance when performance was no longer incentivized. Additional analyses were conducted by adding covariates to this model and evaluating model fit comparisons, as explained below.

To assess whether there was an effect of learning performance on NoGo accuracy, we calculated the proportion optimal choice during the late stage of the learning task (last 60 trials) for each participant for each learning condition (low gain, high gain, low loss, high loss). We computed a mixed-effects model with NoGo accuracy as the dependent variable and fixed effects for proportion optimal choice, stakes, and valence, and a random effect for subject to account for repeated measures. This model tested for a main effect of learning performance (proportion optimal choice) and the interaction terms for learning performance by stakes, learning performance by valence, and learning performance by stakes by valence.

To examine the effect of age on NoGo performance, we conducted a mixed-effects model that included fixed effects for stakes, valence, and continuous linear age, to test for a main effect of age and an age by stakes by valence interaction (the model also included all possible two-way interactions). The model included a random effect for subject to account for repeated measures. We repeated this analysis including additional covariates for Go Accuracy and Go reaction time to control for additional behavioral factors that may have influenced NoGo performance. For example, if Go accuracy was low, this could indicate disengagement from the task, which may artificially inflate NoGo accuracy due to non-responding. Go RT analyses accounted for speed/accuracy tradeoffs (Thorpe et al., 1996), as speeded responses during Go trials may make it more difficult to withhold a response for NoGo trials. For age analyses, post-hoc t-tests with Tukey’s correction (implemented via the lsmeans package in R) were computed for age sub-groups, separated into two-year age bins: 13-14 years, n=19; 15-16 years, n=20; 17-18 years, n=21; 19-20 years, n=21. These post-hoc analyses allowed us to characterize the direction of age interaction effects.

We conducted analyses on self-report valence and arousal ratings to assess whether the subjective pleasantness and intensity associated with winning and losing low and high outcomes during learning varied as a function of valence (gain/loss), stakes (low/high), and age. We conducted mixed-effects models for valence and arousal ratings to test for fixed effects of stakes. valence, continuous linear age, and all interaction terms. The model included a random effect for subject to account for repeated measures. These analyses allowed us to quantify whether subjective experience of the task monetary outcomes was consistent across age, which allowed us to validate that the task incentives were experienced similarly across the age range.

All statistical analyses for behavioral data were conducted in R. Mixed-effects models were computed with the nlme package (Pinheiro et al., 2019). Model fit comparison was calculated by comparing AIC values between models and computing a log-likelihood ratio with the anova function.

### FMRI data acquisition

Participants completed functional MRI scanning in a 3.0 Tesla Siemens Prisma MRI scanner equipped with a 32-channel head coil (Siemens Medical Systems) at the Harvard University Center for Brain Science. Participants viewed the experimental task via a back-projection system. Behavioral tasks were developed and presented with Psychopy version 1.83 (Peirce, 2007). Button responses were recorded with an MRI compatible button box. The presentation of the experimental task was synchronized to fMRI volume acquisition following initial dummy scans collected for magnet stabilization. Functional (T2* weighted) images were collected using an interleaved echo planar sequence using the following parameters: TR 2000ms, TE 35ms, flip angle 80°, 69 axial slices, voxel resolution 2.2×2.2×2.2mm, multi-band acceleration factor of 3. High resolution T1-wighted structural MRI scans were also acquired using a multi-echo multi-planar rapidly acquired gradient echo (MEMPRAGE) sequence and were used to co-register the functional images to a standardized anatomical space (176 sagittal slices, TR 2200ms, TE 1.67ms, flip angle 7°, slice thickness 1mm, voxel size 1×1×1mm).

### FMRI data processing and quality assessment

FMRI data processing and analysis for the Go/NoGo task were conducted with FSL (version 5.0.4) (Smith et al., 2004). Preprocessing was conducted in FSL and implemented through the Lyman pipeline (v.0.0.7, https://github.com/mwaskom), which relies on the Nipype project framework (v. 0.9.2) (Gorgolewski et al., 2011). Standard preprocessing steps included slice-time correction, realignment, coregistration of functional to structural images using bbregister (Greve & Fischl, 2009), non-linear normalization of structural to FSL’s MNI152 template space using ANTS 1.9.x, svn release 891; (Avants et al., 2009), and spatial smoothing with a 6mm Gaussian kernel.

Functional MRI data were carefully evaluated for motion and signal outliers given the negative impact it can have on signal quality and GLM estimates. The following rules were imposed for exclusion of functional data as in our prior work (Insel et al., 2017; Insel & Somerville, 2018). Runs in which more than 10% of TRs were censored for motion (relative motion > 1mm) or outlier signal intensity (exceeded the grand run median by 4.5 median absolute deviations) were excluded from analysis. Runs with a single relative movement exceeding 5 mm were also excluded. In total, 5 participants were excluded from fMRI analyses due to excessive motion.

### Functional activity analysis

Preprocessed BOLD data were submitted to a general linear model (GLM) analysis using film_gls in FSL (Smith et al., 2004) to estimate relevant task effects. Task regressors included temporal onsets for the following events: Block instructions, Go trials, previous high gain cue NoGo trials, previous low gain cue NoGo trials, previous high loss cue NoGo trials, previous low loss cue NoGo trials, Error trials (missed response for Go or false alarm for NoGo). Extensive jittered timings that were incorporated into the experimental design rendered the different trial phases and conditions separately estimable. All task regressors were convolved with the canonical hemodynamic response function. Nuisance regressors included 6-parameter motion estimates, censored frames for deviant signal intensity and excessive motion (> 1mm), whole brain white matter signal, and cerebral spinal fluid signal.

Random-effects group analyses (whole-brain voxelwise t-tests) were conducted to identify task-based changes in functional activity and connectivity, as described below. We conducted whole brain analyses using a series of one-sided t-tests. All whole brain maps were thresholded using whole-brain correction of z>2.3 using FLAME 1+2, as implemented in FSL, resulting in whole-brain threshold of p<0.05 family-wise error (FWE) corrected.

To assess the effects of previous gain learning, fMRI analyses examined the previous high gain cue NoGo > previous low gain cue NoGo contrast. This contrast was selected because these conditions were found to be driving age-related behavioral effects, such that NoGo performance improvements for previous high gain cue NoGo relative to previous low gain cue NoGo trials emerged with age.

To assess age-related differences in functional activity for previously learned gain cues, we added a covariate for mean-centered linear age to the contrast of previous high gain cue NoGo > previous low gain cue NoGo. Age analyses were small-volume corrected using masks for the frontal operculum and striatum (bilateral caudate, putamen, and nucleus accumbens), which were extracted from the Harvard-Oxford Atlas, thesholded at a 10% probability. These regions were selected given their role in motivated cognitive control, and specifically prior findings demonstrating age-related differences in recruitment during Go/NoGo performance (Insel et al., 2017). Small-volume correction was conducted using the easythresh function in FSL. Easythresh is a function that computes statistics, including corrected thresholds and maps, outside of the FEAT environment.

To assess the effects of previous loss learning, fMRI analyses examined the previous high loss cue NoGo > previous low loss cue NoGo contrast. This contrast served as a point of comparison to determine whether the effects of cues associated with previous high stakes learning were valence-selective. To assess age-related differences in functional activity for previously learned loss cues, we added a covariate for mean-centered linear age to the contrast testing previous high loss cue NoGo > previous low loss cue NoGo. Because these age-related analyses in the loss domain were exploratory, maps were subjected to whole brain corrected thresholds.

Cluster peaks are reported in a table for both the whole-group maps and the age covariate maps. We detected clusters that survived correction for the increasing age covariate maps, but not for the decreasing age covariate maps, so only those results are reported below. Region labels are based on the Harvard-Oxford Cortical and Subcortical atlases. Sub-peaks were defined by local maxima (activation peaks) within each cluster using a higher-values-first watershed searching algorithm, as implemented in the lyman pipeline.

### Functional Connectivity Analyses

To assess functional connectivity during the task, we conducted Psychophysiological Interaction (PPI) analyses to examine functional coupling that was enhanced for previously learned high gain relative to previously learned low gain cues during NoGo execution. We focused on this contrast given that the age-related differences in stakes-selective behavior were specific to the gain domain. The caudate was chosen as a seed region given its well-established involvement in value-based learning and action selection (Haber & Knutson, 2010), and the documented anatomical connections between the caudate and prefrontal cortex, which have been shown to support value-guided cognitive control (Ridderinkhof et al., 2004) and mature through adolescence. Caudate seeds were defined by creating anatomical masks from the Harvard Oxford atlas. The PPI interaction regressors were based on timecourses of the left and right caudate seeds extracted using the fslmeants function. Separate PPI analyses were conducted for the left and right seeds. Timeseries were multiplied by the temporal onsets for the previously learned high gain NoGo > previously learned low gain NoGo to create the PPI interaction regressor. PPI analyses incorporated additional timeseries regressors for mean white matter signal and ventricular signal into the General Linear Model, which were added as nuisance regressors to reduce potential noise confounds. To reduce motion-induced confounds which have been shown to distort connectivity profiles, especially in developmental samples (Satterthwaite et al., 2012), nuisance regressors for the six motion parameters for translations and rotations in order were added to the model, and volumes with excess motion were excluded from analysis.

Fixed effects models were conducted at the subject level and then submitted to a random effects analysis to compute the group-level statistics focused on the psychophysiological interaction regressor. Random effects analyses were conducted to identify age effects on connectivity using the mean-centered linear age regressor as a predictor of differential functional connectivity for previously learned high gain versus low gain cues. Two separate age models were computed. The primary age model implemented an age-increasing covariate to identify task-based connectivity that increased across development. The second exploratory age model implemented an age-decreasing covariate to identify task-based connectivity that decreased across development. Finally, a separate whole-brain PPI random effects analysis was conducted to identify brain-behavioral linkages. This model included a covariate for high stakes gain performance enhancements, which was calculated by finding a difference score in NoGo accuracy between previous high gain and previous low gain cues. This model identifed task-based connectivity that increased with increasing stakes-based performance enhancements.

## Results

### Effects of stakes and valence on learning performance

Proportion optimal choice was computed as an index of learning performance, quantifying how frequently individuals chose the cue that received the favorable outcome on the majority of trials. Analyses of proportion optimal choice data revealed that there was a main effect of age, F(1,79)=5.44, p=0.02, whereby the proportion of the time participants selected the optimal choice across all trial types increased with age. There was also a main effect of stakes, F(1,237)=4.53, p=0.03, whereby learning performance improved for high stakes incentives. These main effects were qualified by a significant age by stakes (low/high) by valence (gain/loss) interaction, F(1,237)=7.11, p=0.008. These findings indicated that there were age-related differences in whether optimal choice differed between low and high stakes conditions for trials in which participants were learning about gains versus avoiding losses. Across adolescence, increasing age was associated with emerging improvements in learning from high relative to low stakes outcomes in the gain condition. However, in the loss domain, performance did not differ by stakes with age.

### Effects of previously learned value-associations on subsequent cognitive control

In this task, participants first learned associations between cues that could yield low and high gain and loss incentives during a probabilistic learning task, and then they completed a Go/NoGo cognitive control task in which they were instructed to withhold responding to these cues. To measure the effect of previous learning on Go/NoGo performance, we quantified NoGo accuracy for each condition based on the factorial crossing of stakes (high, low) and valence (gain, loss). Results revealed a main effect of stakes, F(1,236)=10.32, p=0.002, demonstrating that NoGo performance was better to previous high stakes cues (M=0.77, SD=0.16) compared to previous low stakes cues (M=0.74, SD=0.17). The main effects of valence, F(1,236)=0.83, p=0.36, and the stakes by valence interaction, F(1,236)=2.32, p=0.13, were not significant.

We next added previous learning performance for each condition to the model, which was indexed by proportion optimal choice, a behavioral measure that indicates the proportion of trials on which participants made the choice that most often yielded the favorable outcome. The main effect of optimal choice was not significant, F(1,236)=3.35, p=0.07. The interactions of optimal choice by stakes, F(1,236)=0.34, p=0.56, optimal choice by valence, F(1,236)=0.90, p=0.34, and optimal choice by stakes by valence, F(1,236)=1.14, p=0.29, were not significant.

We conducted further analyses to assess whether age-related differences in learning performance influenced the observed age-related interactions for NoGo accuracy. To do this, we repeated the existing analyses for NoGo performance and tested a four-way interaction between age, optimal choice, stakes, and valence and all two-way and three-way interactions. When including these interaction terms, the age by valence by stakes interaction remained significant for NoGo accuracy, F(1,229)=5.14, p=0.02. If learning performance influenced the age-related interactions we observed for NoGo accuracy, then we would expect that the 4-way interaction term would be significant. However, the stakes by valence by age by optimal choice interaction was not significant, F(1,229)=0.20, p=0.65.

Model fit comparison revealed that adding optimal choice to the model did not significantly improve model fit (AIC model without learning performance=-436.20, AIC model with learning performance=-433.31, log-likelihood ratio=5.23, p=0.27). Therefore, optimal choice performance was not included in subsequent analyses.

### Cognitive control facilitation from prior high stakes gains emerges with age

Key analyses assessed whether the effect of previously learned value-associations differentially influence cognitive control performance across adolescence as a function of age. To test this effect on NoGo accuracy, we added continuous linear age to the model that included factors for stakes and valence. Adding the age term to the model significantly improved model fit (AIC model without age=-436.20, AIC model with age=-438.35, log-likelihood ratio=10.15, p=0.04). There was a significant main effect of age, F(1,79)=5.87, p=0.02, revealing that NoGo accuracy improved with age. The main effect of stakes was significant as in the previous model, F(1,237)=10.37, p=0.002.

Notably, there was a significant age by stakes by valence interaction, F(1,237)=4.18, p=0.04. To interrogate the direction of this interaction, we performed post-hoc t-tests between low and high stakes within the gain and loss conditions for two-year binned age groups. There was a significant improvement for high stakes gain cues relative to low stakes gain cues in the 19-20 year old group (p=0.006), but this difference was not significant for the other age groups (Table 1). The difference between NoGo performance for low stakes loss cues and high stakes loss cues did not differ for any of the age groups (Table 1). Together, these results reveal that NoGo accuracy improvements for high stakes relative to low stakes cues emerged with age for previous gain cues but not for previous loss cues.

**Table 1:** Post-hoc comparisons for NoGo Accuracy.

| **Table 1: Post-hoc comparisons for NoGo Accuracy** |  |  |  |  |
| --- | --- | --- | --- | --- |
| High vs. Low Contrast | 13-14 | 15-16 | 17-18 | 19-20 |
| Previous Gain Cue | t=0.29, p=0.77 | t=1.59, p=0.12 | t=1.73, p=0.09 | t=2.82, p=0.006 |
| Previous Loss Cue | t=1.44, p=0.16 | t=0.28, p=0.78 | t=1.27, p=0.21 | t=-0.64, p=0.53 |
| *p-values corrected for multiple comparisons with Tukey's HSD method* |  |  |  |  |

### Control analyses for age-related performance improvements

We conducted control analyses to account for other Go/NoGo performance variables that may have influenced NoGo performance, including Go accuracy and Go reaction time (RT). For example, if an individual was responding faster to Go trials, this could worsen performance due to speed/accuracy tradeoffs, whereas if an individual was failing to respond to Go cues, this could artificially enhance NoGo accuracy due to disengagement from the task. To test this, we added Go accuracy and Go RT to the model testing the age by stakes by valence interaction reported above. Adding Go accuracy did not significantly improve model fit (AIC model without Go accuracy=-436.20, AIC model with Go accuracy=-436.63, log-likelihood ratio=10.42, p=0.06), and the main effect of Go accuracy was not significant, F(1,236)=0.16, p=0.69. When adding Go accuracy to the model, the main effect of age F(1,79)=5.89, p=0.02, stakes F(1,236)=10.33, p=0.001, and the age by stakes by valence interaction F(1,236)=4.13, p=0.043, remained significant.

Adding Go RT did significantly improve model fit (AIC model without Go RT=-436.20, AIC model with Go RT=-439.63, log-likelihood ratio=13.43, p=0.02). However the main effect of Go RT was not significant, F(1,236)=2.80, p=0.10. When controlling for Go RT, the main effects of age F(1,79)=6.63, p=0.01, stakes F(1,236)=10.05, p=0.002, and the age by stakes by valence interaction F(1,236)=3.97, p=0.047, remained significant.

### Subjective value ratings

Participants completed self report pleasantness and arousal ratings to quantify subjective value of the monetary outcomes encountered in the learning task. These ratings served as a manipulation check to verify that participants differentially valued the monetary outcomes in the learning task. These ratings also allowed us to complete age-related control analyses to verify that subjective valuation of the monetary outcomes did not systematically vary with age.

For arousal ratings, there was a main effect of stakes F(1,71)=131.03, p<0.001, whereby participants rated high stakes outcomes to be significantly more arousing than low stakes outcomes. There was no main effect of valence on arousal ratings F(1,71)=0.07, p=0.79, such that arousal ratings were comparable for gain and loss conditions. In addition, there was no main effect of age on arousal ratings, F(1,71)=0, p=0.99 and no interactions with age (see Table 2), indicating that subjective arousal was comparable across the age range.

**Table 2:** Hedonic Experience Ratings ANOVA table.

| **Table 2: Hedonic Experience Ratings ANOVA table** |  |  |
| --- | --- | --- |
|  | Arousal Ratings | Pleasantness Ratings |
| Stakes | $F(1,71)=131.03$, $p<0.001$ | $F(1,71)=8.42$, $p=0.005$ |
| Valence | $F(1,71)=0.07$, $p=0.79$ | $F(1,71)=406.6$, $p<0.001$ |
| Age | $F(1,71)=0.00$, $p=0.99$ | $F(1,71)=0.73$, $p=0.40$ |
| Stakes * Valence | $F(1,71)=1.86$, $p=0.18$ | $F(1,71)=157.68$, $p<0.001$ |
| Stakes * Age | $F(1,71)=2.01$, $p=0.16$ | $F(1,71)=1.35$, $p=0.25$ |
| Valence * Age | $F(1,71)=0.19$, $p=0.67$ | $F(1,71)=1.23$, $p=0.27$ |
| Stakes * Valence * Age | $F(1,71)=0.12$, $p=0.73$ | $F(1,71)=0.00$, $p=0.98$ |

For pleasantness ratings, there was a main effect of stakes, F(1,71)=8.42, p=0.005 and valence, F(1,71)=406.6, p<0.001, which was qualified by a stakes by valence interaction F(1,71)=157.68, p<0.001. This demonstrates that participants found high stakes gains to be significantly more positive than low stakes gains t(72)=4.86, p<0.001, whereas high stakes losses were significantly more negative than low stakes losses t(72)=-6.69, p<0.001. Importantly, there was no main effect of age on valence ratings F(1,71)=0.73, p=0.40, and no interactions with age were significant (see Table 2). Therefore, subjective ratings of the monetary incentives were consistent with age.

### FMRI results

#### Functional activity analyses

Neuroimaging analyses for cues previously associated with gain outcomes focused on comparisons of functional recruitment for previously learned high stakes relative to low stakes gain cues during NoGo performance, given that the behavioral effects reported above revealed that the improvements to previously learned high stakes cues that emerged with age were selective to the gain domain. To test for stakes effects on previous gain cue NoGo trials, we first computed a whole-brain contrast for high gain NoGo cues versus low gain NoGo cues, isolating trials when participants successfully withheld a response. This group contrast revealed that there was enhanced activity for high stakes gain cues relative to low stakes gain cues in the lateral occipital cortex, insula, occipital pole, and postcentral gyrus (Figure 3a, Table 3).

**Table 3.** Cluster table for previous high gain versus low gain NoGo contrast, FWE p<0.05.

| **Previous High Stakes Gain > Previous Low Stakes Gain NoGo** |  |  |  |  |  |
| --- | --- | --- | --- | --- | --- |
| **Group Mean** |  |  |  |  |  |
| **Region** | **x** | **y** | **z** | **k** | **z-stat** |
| **Lateral Occipital Cortex** | 34 | -86 | -12 | 5898 | 4.93 |
| Superior Temporal Gyrus | 66 | -30 | 10 |  | 4.33 |
| Temporal Pole | 62 | -8 | -2 |  | 4.09 |
| Insular Cortex | 40 | -16 | 4 |  | 3.11 |
| Occipital Pole | 20 | -100 | 14 |  | 2.6 |
| **Occipital Pole** | -12 | -94 | -14 | 1802 | 4.64 |
| Lateral Occipital Cortex | -48 | -70 | 0 |  | 3.58 |
| Temporal Occipital Fusiform Cortex | -26 | -48 | -18 |  | 2.96 |
| **Postcentral Gyrus** | -52 | -8 | 20 | 1689 | 4.03 |
| Planum Temporale | -44 | -40 | 14 |  | 3.15 |
| **Increasing With Age\*** |  |  |  |  |  |
| **Region** | **x** | **y** | **z** | **k** | **z-stat** |
| **Frontal Operculum Cortex** | 48 | 18 | 0 | 28 | 3.21 |
| Frontal Operculum Cortex | 48 | 10 | 2 |  | 2.87 |
| **Caudate (left)** | -4 | 2 | 6 | 53 | 2.74 |
| **Caudate (right)** | 8 | 4 | 10 | 27 | 2.74 |
| *\*No regions showed decreasing activity with increasing age* |  |  |  |  |  |

**Figure 3.**
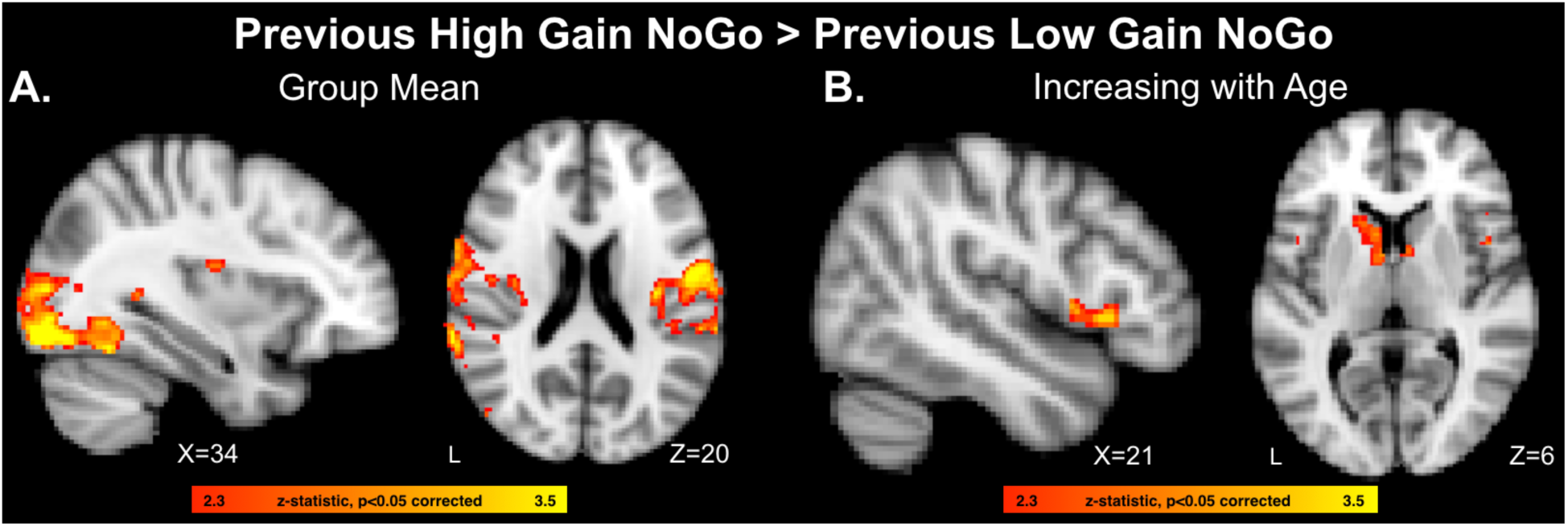
A) Whole-brain analysis identifying neural regions exhibiting enhanced recruitment for previously learned high stakes gain NoGo cues relative to previous low stakes gain NoGo cues in the full sample. B) Neural regions demonstrating linear age-related increases in recruitment for the previously learned high stakes gain NoGo > previous low stakes gain NoGo contrast.

To assess the effects of age on functional recruitment for previous gain cues, we conducted further analyses for the high gain cue NoGo > low gain cue NoGo contrast that included a covariate for continuous linear age. Because we had specific hypotheses about the role of corticostriatal circuitry in value-guided cognitive control (Botvinick & Braver, 2015; Davidow et al., 2018a), we constrained this analysis to interrogate voxels within anatomical masks comprising the bilateral vlPFC (frontal operculum) and bilateral ventral and dorsal striatum (caudate, putamen, and nucleus accumbens). These regions were chosen based on prior work which found enhanced recruitment of these systems during incentivized control in the same 13 to 20 year old age range tested in the present study (Insel et al., 2017). This age covariate analysis identified increased activity for previously learned high gain relative to low gain NoGo cues that increased with age across adolescence in the caudate and in the frontal operculum, extending to the inferior frontal gyrus (Figure 3b, Table 3).

Neuroimaging analyses for cues previously associated with loss outcomes focused on comparisons of functional recruitment for previously learned high stakes relative to low stakes loss cues during NoGo performance. These analyses of the previous loss cues serve as a relative comparison for the previous gain cues to test for valence specificity in the effects of previously learned high stakes on subsequent functional recruitment during NoGo performance. To test for stakes effects on previous loss cue NoGo trials, we first computed a whole-brain contrast for high loss NoGo cues versus low loss NoGo cues, isolating trials when participants successfully withheld a response. This group contrast revealed that there was enhanced activity for high stakes loss cues relative to low stakes loss cues in the occipital pole, lingual gyrus, lateral occipital cortex, and inferior temporal gyrus (Table 4).

**Table 4.** Cluster table for previous high loss versus low loss NoGo contrast, FWE p<0.05.

| **Previous High Stakes Loss > Previous Low Stakes Loss NoGo Cues** |  |  |  |  |  |
| --- | --- | --- | --- | --- | --- |
| **Group Mean** |  |  |  |  |  |
| **Region** | **x** | **y** | **z** | **k** | **z-stat** |
| **Occipital Pole** | 2 | -94 | 22 |  | 5.57 |
| Lingual Gyrus | -6 | -86 | -20 |  | 5.33 |
| Lateral Occipital Cortex | 44 | -70 | -26 |  | 4.67 |
| Occipital Pole | 22 | -96 | -4 |  | 4.36 |
| Inferior Temporal Gyrus | -50 | -56 | -24 |  | 4.11 |
| **Increasing With Age\*** |  |  |  |  |  |
| **Region** | **x** | **y** | **z** | **k** | **z-stat** |
| **Precuneus** | 2 | -74 | 44 | 559 | 3.59 |
| Precuneus | 2 | -48 | 60 |  | 2.72 |
| *\*No regions showed decreasing activity with increasing age* |  |  |  |  |  |

To assess the effects of age on functional recruitment for previous loss cues, we conducted further analyses for the high loss cue NoGo > low loss cue NoGo contrast that included a covariate for continuous linear age. This age covariate analysis identified increased activity for previously learned high loss relative to low loss NoGo cues that increased with age across adolescence in the precuneus (Table 4).

### Functional connectivity analyses

We conducted PPI analyses to examine how previously learned high stakes gain cues modulate functional coupling with the striatum. The left and right caudate were used as seed regions, given the its role in reward learning and action selection, and the PPI analysis targeted the contrast for previously learned high gain cues > previously learned low gain cues. There were no significant clusters that survived whole-brain corrected thresholding for the full group average analysis. Our primary question, however, was whether differences in connectivity changed with age. With decreasing age, there was enhanced coupling between the caudate, nucleus accumbens, and ventromedial prefrontal cortex for previously learned high gain relative to previously learned low gain cues (left caudate seed, Figure 4A, Table 5). With increasing age, there was enhanced coupling between the caudate and dorsolateral prefrontal cortex (right caudate seed, Figure 4B, Table 5).

**Table 5.**
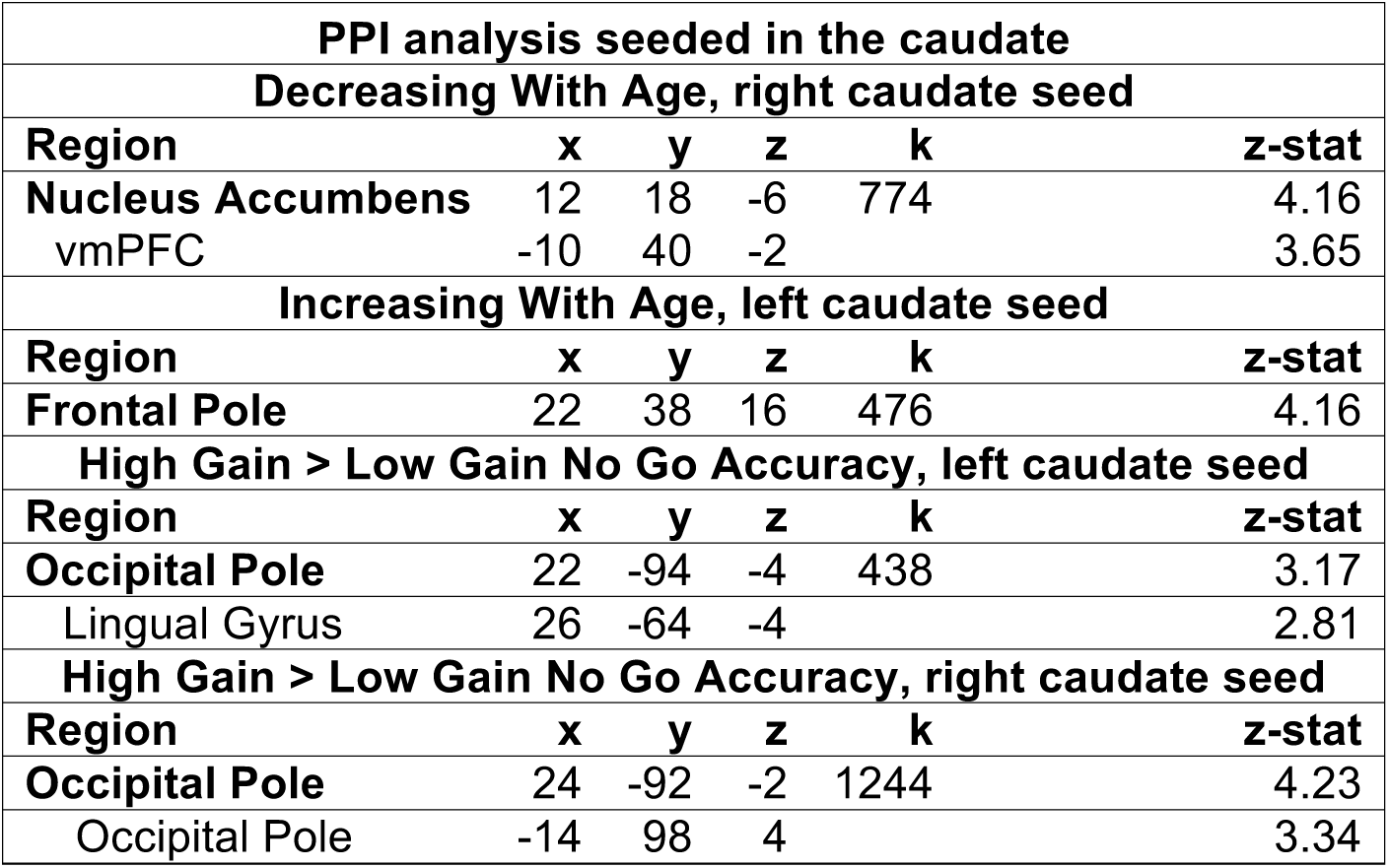
Cluster table for PPI analysis, FWE p<0.05.

| **PPI analysis seeded in the caudate** |  |  |  |  |  |
| --- | --- | --- | --- | --- | --- |
| **Decreasing With Age, right caudate seed** |  |  |  |  |  |
| **Region** | **x** | **y** | **z** | **k** | **z-stat** |
| **Nucleus Accumbens** | 12 | 18 | -6 | 774 | 4.16 |
| vmPFC | -10 | 40 | -2 |  | 3.65 |
| **Increasing With Age, left caudate seed** |  |  |  |  |  |
| **Region** | **x** | **y** | **z** | **k** | **z-stat** |
| **Frontal Pole** | 22 | 38 | 16 | 476 | 4.16 |
| **High Gain > Low Gain No Go Accuracy, left caudate seed** |  |  |  |  |  |
| **Region** | **x** | **y** | **z** | **k** | **z-stat** |
| **Occipital Pole** | 22 | -94 | -4 | 438 | 3.17 |
| Lingual Gyrus | 26 | -64 | -4 |  | 2.81 |
| **High Gain > Low Gain No Go Accuracy, right caudate seed** |  |  |  |  |  |
| **Region** | **x** | **y** | **z** | **k** | **z-stat** |
| **Occipital Pole** | 24 | -92 | -2 | 1244 | 4.23 |
| Occipital Pole | -14 | 98 | 4 |  | 3.34 |

**Figure 4.**
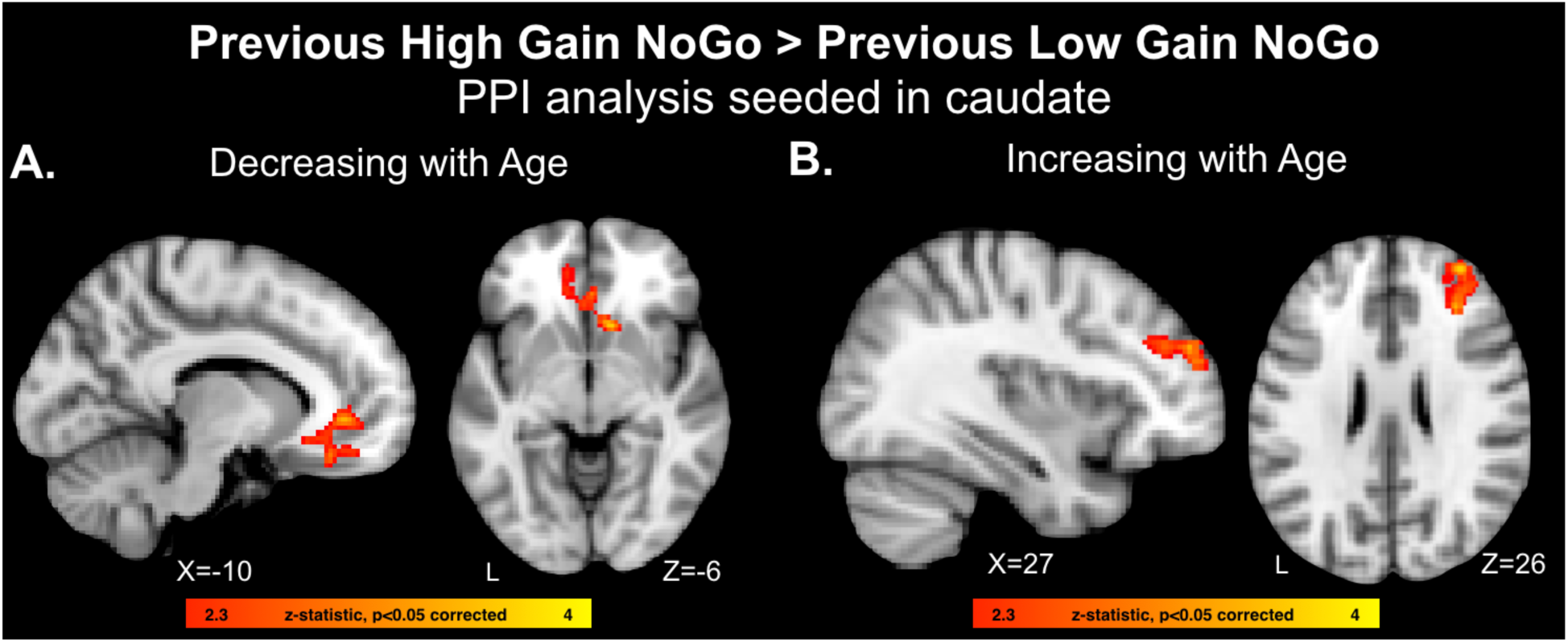
A) PPI analysis identifying age-related decreases (stronger in younger adolescents) in functional coupling with the caudate for previously learned high stakes gain NoGo cues relative to previous low stakes gain NoGo cues that decreased with age. B) PPI analysis identifying age-related increases (stronger in older adolescents) in functional coupling with the caudate for previously learned high stakes gain NoGo cues relative to previous low stakes gain NoGo cues that increased with age.

To assess whether functional connectivity was related to behavioral differences on the task, we conducted a PPI analysis to identify loci in which greater coupling with the caudate was associated with enhanced NoGo accuracy for high stakes gain cues relative to low stakes gain cues. For both the right and left caudate seeds, improved accuracy for high stakes gain cues was associated with enhanced coupling between the caudate and visual cortex (Table 5) during previous high relative to previous low gain NoGo cues.

## Discussion

The present study examined how previously learned value associations influence the subsequent execution of cognitive control differentially with age across adolescence, and interrogated the corresponding neurodevelopmental processes. We first trained cue-incentive associations for low and high stakes gain and loss outcomes through a reinforcement learning task, and then tested how these learned value associations influenced the ability to withhold a motor response when these cues were presented in a subsequent Go/NoGo task and performance was no longer incentivized. We compared behavioral and neural responses in participants aged 13 to 20 who completed the post-learning Go/NoGo task while undergoing functional neuroimaging.

Behavioral analyses revealed that there was an emerging improvement in NoGo accuracy with age for previously learned high-gain relative to low-gain cues, but not for previous loss cues. Specifically, older adolescents 19-20 years old improved NoGo performance for previously learned high relative to low gain cues, but they did not differentiate performance by high relative to low loss cues. Neuroimaging analyses assessed differences in functional recruitment for previously learned high stakes gain relative to low stakes gain NoGo cues. Age analyses revealed that, with age, participants exhibited increased recruitment of the ventrolateral prefrontal cortex (vlPFC) and caudate for previous high stakes gain cues relative to previous low stakes gain cues. Further, for previous high gain relative to previous low gain cues, coupling between the caudate and dorsolateral prefrontal cortex (dlPFC) increased with age, whereas coupling between the caudate and ventromedial prefrontal cortex (vmPFC) decreased with age. Together, these findings reveal that previously learned value-associations acquired through gain learning facilitate subsequent cognitive control in older adolescents, and this facilitation emerges with age during adolescence. Moreover, the late adolescent emergence of improved cognitive control over high-value gain cues was paralleled by age-related enhancements in functional recruitment and connectivity within corticostriatal circuitry. These findings may reflect a late-emerging tendency to integrate value associations with cognitive control demands to adjust performance in a value-driven fashion.

The key question of the present study was whether previously learned value associations, trained through probabilistic reinforcement learning, would transfer beyond the learning environment to influence the execution of cognitive control when incentives are no longer at stake. To test this, participants first learned cue-incentive associations through reinforcement learning, when selecting a cue was associated with low stakes or high stakes gain or loss outcomes. Next, participants were instructed to withhold responding to the previously learned cues in a Go/NoGo task, when performance was no longer incentivized. Consistent with prior developmental research, we found that the overall ability to execute cognitive control by withholding a motor response improved with age (Bunge & Wright, 2007; Crone & Steinbeis, 2017; Luna et al., 2004). However, this main effect of age was qualified by an interaction with previously learned stakes (low/high) and valence (gain/loss) associations. Specifically, with age across adolescence, individuals were more likely to improve NoGo accuracy to cues previously associated with high stakes gain outcomes relative to cues previously associated with low stakes gain outcomes. However, NoGo accuracy for previous loss cues did not vary by stakes with age. Importantly, this age-related pattern in post-learning cognitive control performance could not be explained by individual differences in learning performance or age-related differences in subjective value of the incentives.

The high-value gain facilitation effect that emerged in late adolescence mirrors prior research in young adult samples. For example, prior studies have demonstrated that when cues are first associated with incentives and reintroduced in a subsequent cognitive task, adults typically improve cognitive performance when encountering cues previously paired with high value outcomes, even when performance on this secondary task is no longer incentivized (Krebs & Woldorff, 2017). Theoretical models based on the young adult literature posit that previously acquired high-value associations may modulate visual attention, which can orient goal-directed action selection (Padmala & Pessoa, 2011; Pessoa & Engelmann, 2010) along with value-selective recruitment of visual cortex, which is believed to orient attention and perceptual discrimination to sharpen cognitive performance (Della Libera & Chelazzi, 2006; Serences, 2008). Results from the present study are consistent with this idea, as we observed enhanced recruitment of visual cortex for previously learned high gain relative to low gain cues across the full group. Moreover, better NoGo performance for the previous high gain cues relative to the previous low gain cues was associated with enhanced functional coupling between the caudate and visual cortex. This suggests that value-selective communication between the striatum and visual cortex could improve cognitive control performance by enhancing bottom-up visual attention, which may aid in visual discrimination and facilitate action selection.

The present results revealed that behavioral facilitation from previously learned high gain value cues emerged over the course of adolescence in parallel with value-selective upregulation of corticostriatal regions. Specifically, we found that with increasing age, participants enhanced recruitment of the caudate and vlPFC for NoGo cues associated with prior high gain relative to low gain outcomes during the previous learning task. However, this age-related pattern was not observed when comparing responses between previous high loss and previous low loss cues, which detected age-related differences in the precuneus. This suggests that the value-selective enhancements of corticostriatal recruitment, which emerged with age, were specific to the gain domain. The caudate is often implicated in representing the value of a stimulus and coordinating motor output, and the vlPFC is thought to orchestrate action selection and execute cognitive control (Ridderinkhof et al., 2004). Notably, these regions share connections within a broad and distributed neural circuit (Haber & Knutson, 2010). Within this network, functional recruitment of the dorsal striatum and lateral PFC, and connectivity between these regions, increases with age alongside age-related improvements in motor control (Church et al., 2017; Somerville et al., 2011; Vink et al., 2014). Further, as individuals age across adolescence, they are more likely to enhance recruitment of this network while engaging in more complex cognitive control strategies (Vink et al., 2014). Therefore, individuals may be more likely to use value cues to adjust control performance with age when functional maturation of these systems begins to stabilize and cognitive control execution becomes more adaptive (Davidow et al., 2018a).

Functional connectivity analyses identified preferential responses to previously learned high gain relative to previously learned low gain cues that changed with age. These results revealed distinct shifts in corticostriatal connectivity, such that younger adolescents expressed enhanced coupling between the caudate, ventral striatum, and vmPFC for previously learned high gain cues relative to low gain cues. However, with increasing age, older adolescents expressed enhanced coupling between the caudate and dlPFC. These findings suggest that for previously learned high value cues, connectivity with the striatum undergoes a medial to lateral shift with age. The vmPFC has been consistently implicated in coding subjective value and guiding choice behavior (Roy et al., 2012), whereas the dlPFC is more commonly implicated in the execution of higher order cognitive control (Badre, 2008). Thus, while younger adolescents may exhibit neural signatures of value-based learning that persist beyond the learning context, these associations may not confer behavioral benefits. In contrast, younger adolescents improved performance for previously learned high value cues in conjunction with enhanced caudate-dlPFC connectivity. Together, these results reveal that age-related shifts in value-guided functional connectivity may account for the late emerging improvements in cognitive control over previously incentivized high value cues.

Taken together, these findings suggest that across adolescence, the emerging tendency to integrate previously learned value-associations with subsequent cognitive control demands may be subserved by selective recruitment of corticostriatal systems. These findings build on a neurodevelopmental framework that posits that during adolescence, the ability to integrate value-related information with cognitive control demands emerges with age when cognitive control abilities reach a point of stable maturation (Davidow et al., 2018a). The present findings extend this framework by identifying a similar age-related pattern for performance that is no longer incentivized, but which contains cues previously associated with valued outcomes. Therefore, these value associations can confer benefits in cognitive control during late adolescence, even when these cues are no longer incentivized.

The present findings, which demonstrate the late adolescent emergence of value-based performance facilitation, diverge from prior work implementing a similar experimental approach in a developmental sample with an overlapping age range. Specifically, Davidow and colleagues (2018b) found that when value associations are first trained through a speeded instrumental reward task (monetary-incentive delay task), which requires participants to respond increasingly quickly to a single cue to earn monetary rewards, this value history actually intrudes on subsequent cognitive control performance. Notably, this value-related intrusion emerged during adolescence and persisted into young adulthood. Therefore, while the effect on performance was opposite of what we observed in the present study, the age-related pattern of when value history influences subsequent performance is similar in the present study.

Why would value cues facilitate performance in late adolescence in the present study, but hinder performance in this prior work? A recent framework of motivation/cognition interactions suggests that reward value-associations help when trained value cues are congruent with task demands in the subsequent cognitive control test, but these associations can hinder performance when they are incongruent and distract attention away from the central task (Krebs et al., 2011; Krebs & Woldorff, 2017). Therefore, the motor components of how the value-associations are first trained can influence their subsequent interactions with control implementation. For example, Davidow and colleagues trained value associations through response speeding, which may bind value associations with immediate motor actions. When these cues are presented in a new context in which the participant must withhold a response, this motor/incentive association makes response inhibition difficult because it is incongruent with the learned association. Future studies will need to explicitly test this proposed dissociation.

The present study first trained value associations using a reinforcement learning task, in which individuals could deliberate for up to three seconds when making a choice, and there was no inherent utility to response speeding. Therefore, this training through learning may link attentional features with valued outcomes more than motoric-based associations, which could, in turn, benefit subsequent performance. Thus, we speculate that the facilitation observed in the present study may stem from high value cues orienting visual attention towards salient and task-relevant cues. Another possibility is that high value associations enhance context monitoring (Chatham et al., 2012; Chevalier et al., 2014; Winter & Sheridan, 2014), a cognitive control process that involves selectively attending to the environment for relevant cues to determine the contextually appropriate action to select. Maturing memory systems may also account for the age-related differences we observed in value-guided cognitive control. Future work is needed isolate these distinct cognitive mechanisms to better understand how key components of value-based transfer shift during development.

In summary, the present study tested how previously learned cue-incentive associations differentially influence subsequent cognitive control across age during adolescence. For behavioral performance, no-go accuracy increased with age for previously learned high-gain relative to low-gain cues, but not for previous loss cues, and this beneficial effect of high-gain cues emerged in late adolescence. This suggests that for older adolescents, high-value associations transfer across cognitive domains and improve later goal directed behavior. These beneficial effects of high value gain cues may result from enhanced attentional modulation or increased context monitoring, processes which allow an individual to optimize action selection in the moment. Neuroimaging analyses demonstrated that with increasing age, individuals were more likely to enhance recruitment of the dorsal striatum and ventrolateral prefrontal cortex to previously learned high gain cues relative to previously learned low gain cues. Further, older adolescents enhanced functional connectivity between the dorsal striatum and dorsolateral prefrontal cortex when inhibiting responses to previously learned high gain cues. Together, these findings reveal that the persistent beneficial effects of learned high value cue-incentive associations on cognitive control performance emerge in late adolescence in parallel with value-selective recruitment of corticostriatal systems that guide goal-directed behavior.

## Acknowledgements

We thank Barbara Braams, Juliet Davidow, Gina Falcone, Laurel Kordyban, Erik Nook, Katherine Powers, Alexandra Rodman, Maheen Shermohammed, and Constanza Vidal Bustamante for assistance with data collection. We thank Tammy Moran, Ross Mair, and the Center for Brain Science at Harvard University for technical assistance during data collection. We thank Erik Kastman, and Mahalia Prater Fahey for help with data analysis. This research was supported in part by the National Science Foundation (Graduate Research Fellowship DGE 1144152), Harvard Catalyst (Pilot Grant UL1 TR001102), the National Institutes of Health (NIH shared instrumentation grant S10OD020039), and the Sackler Scholar Programme in Psychobiology.

## Declarations of interest

None

